# Single-cell transcriptional dynamics and origins of neuronal diversity in the developing mouse neocortex

**DOI:** 10.1101/409458

**Authors:** L. Telley, G. Agirman, J. Prados, S. Fièvre, P. Oberst, I. Vitali, L. Nguyen, A. Dayer, D. Jabaudon

**Author notes:** equally contributed to this work (L. Telley, G. Agirman).

## Abstract

During cortical development, distinct subtypes of glutamatergic neurons are sequentially born and differentiate from dynamic populations of progenitors. The neurogenic competence of these progenitors progresses as corticogenesis proceeds; likewise, newborn neurons transit through sequential states as they differentiate. Here, we trace the developmental transcriptional trajectories of successive generations of apical progenitors (APs) and isochronic cohorts of their daughter neurons using parallel single-cell RNA sequencing between embryonic day (E) 12 and E15 in the mouse cerebral cortex. Our results identify the birthdate- and differentiation stage-related transcriptional dynamics at play during corticogenesis. As corticogenesis proceeds, APs transit through embryonic age-dependent molecular states, which are transmitted to their progeny to generate successive initial daughter cell identities. In neurons, essentially conserved post-mitotic differentiation programs are applied onto these distinct AP-derived ground states, allowing temporally-regulated sequential emergence of specialized neuronal cell types. Molecular temporal patterning of sequentially-born daughter neurons by their respective mother cell thus underlies emergence of neuronal diversity in the neocortex.

**One Sentence Summary:** During corticogenesis, temporally dynamic molecular birthmarks are transmitted from progenitors to their post-mitotic progeny to generate neuronal diversity.

---

The cerebral cortex is a cellularly heterogeneous structure, whose neuronal circuits underlie high-order cognitive and sensorimotor information processing. During embryogenesis, distinct subtypes of glutamatergic neurons are sequentially born and differentiate from populations of progenitors located in the germinal zones below the cortex (Jabaudon 2017; Florio & Huttner 2014). The aggregate neurogenic competence of ventricular zone progenitors (*i.e.* apical progenitors, APs) progresses as corticogenesis proceeds (Govindan & Jabaudon 2017; Okamoto et al. 2016; Gao et al. 2014, Guo et al. 2013; Gaspard et al. 2007); likewise, newborn neurons transit through sequential transcriptional states as they differentiate (Zahr et al. 2018; Telley et al. 2016; Azim et al. 2009). Although the single-cell transcriptional diversity of the neocortex is increasingly well characterized (Saunders et al. 2018; Zeisel et al. 2018; Kageyama et al. 2018; Nowakowski et al. 2017; Tasic et al. 2016; Zeisel et al. 2015), little is yet known about the molecular processes driving either the progression of AP competence, or the specific differentiation of daughter neurons born from these progenitors at sequential embryonic ages.

To address these questions, we used FlashTag (FT), a high temporal resolution method to pulse-label APs and their daughter neurons (Telley et al. 2016; Govindan et al. 2018), to trace the transcriptional trajectories of successive generations of APs and isochronic cohorts of their daughter neurons between embryonic day (E) 12 and E15. This corresponds to the period during which APs successively generate layer (L) 6, L5, L4 and L2/3 neurons (Jabaudon 2017). Following microdissection of the putative somatosensory cortex, we collected FT^+^ cells by FACS after either 1 h, as APs are still dividing, 24 h, as daughter cells are transiting through the subventricular zone, or 96 h (*i.e.* four days), once daughter neurons have entered the cortical plate (Fig. 1A and fig. S1, A and B) (Telley et al. 2016; Govindan et al. 2018). We performed single-cell RNA sequencing at each of these 3 differentiation stages and 4 embryonic ages (E12, E13, E14, and E15), which yielded a total of 2,756 quality-controlled cells across 12 conditions for analysis (fig. S1, C and D, and Methods).

**Fig. 1.**
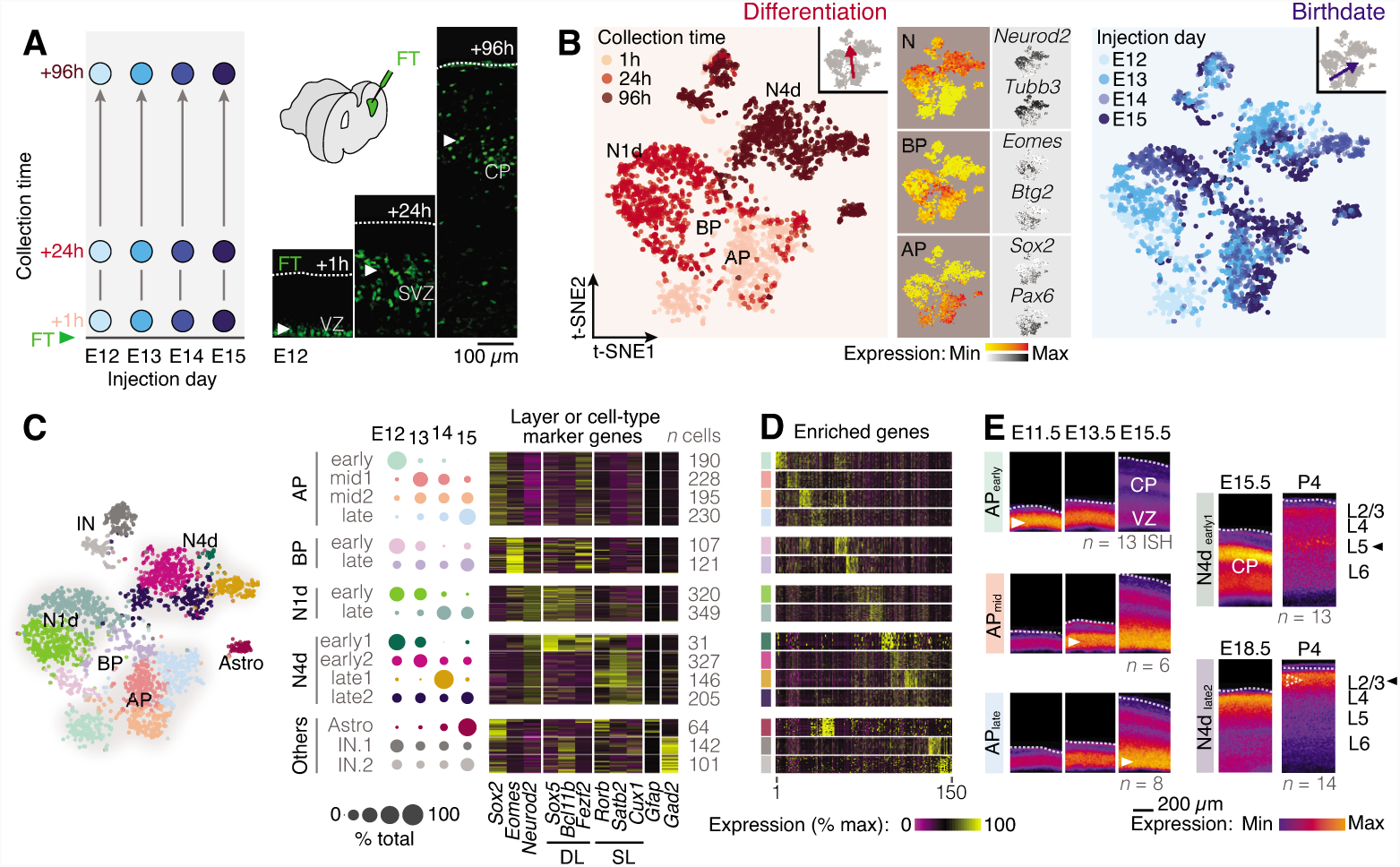
Birthdate and differentiation stage-related cellular diversity in the developing neocortex. (A) Schematic illustration of the experimental procedure. M-phase APs were labeled by FT injection performed at either E12, E13, E14 or E15 and isochronic cohorts of APs and daughter cells were collected either 1, 24 or 96 hours later. (B) t-SNE representation of the single cell RNA sequencing dataset revealing the transcriptional organization of the cells according to the time at which they were collected (i.e. their differentiation status) and the day on which the injection was performed (i.e. their birthdate). APs, BPs and Ns can be distinguished by their combinatorial expression of key marker genes (n = 20 transcripts). (C) Cluster analysis reveals transcriptomically distinct and temporally dynamic cellular clusters. Cluster nomenclature reflects prevalence of the cluster at a given embryonic age (early: E12/E13, late: E14/E15). Cells in these clusters express classical layer and cell-type marker genes in accordance with their birthdate and differentiation status. (D) Expression of the top 150 most highly variable genes highlights cluster diversity. (E) Spatio-temporal expression of cluster-specific transcripts with in situ hybridization. (ISH), from the Allen Developing Mouse Brain Atlas. Color-coded images represent the average expression for representative transcripts (see also Supplementary Figs 3 and 4). Abbreviations: AP: apical progenitors, BP: basal progenitors, Astro: astrocyte, IN: interneurons, N1d: 1-day-old neurons, N4d: 4-day-old neurons, VZ: ventricular zone, SVZ: subventricular zone, CP: cortical plate, DL: deep layer, SL: superficial layer.

Analysis of cellular transcriptional identities using t-SNE dimensionality reduction revealed two main axes of organization: a differentiation (*i.e.* collection time) axis and a birthdate (*i.e.* injection day) axis (Fig. 1B). Along the differentiation axis (Fig. 1B, left), 1 h-, 1-day- and 4-day-old cells were organized into clusters which corresponded to (1) APs, (2) basal progenitors (BPs) and 1-day-old neurons (N1d), and (3) 4-day-old neurons (N4d), as indicated by the combined expression of type-specific markers (Telley et al. 2016). Cells born at successive times of corticogenesis were organized perpendicularly to this differentiation axis, forming a birthdate axis (Fig. 1B, right). This chronotopic map was particularly apparent for APs and 1-day-old daughter cells, but less striking in 4-day-old neurons, suggesting that the salience of birthdate-related transcriptional features decreases with differentiation. Together, these data reveal two orthogonal axes of transcriptional organization: a differentiation axis, corresponding to the birth and maturation of daughter neurons, and a birthdate axis, corresponding to the temporal progression in AP transcriptional states at sequential embryonic ages. These two cardinal processes are the major source of transcriptional diversity in the developing neocortex.

We used a graph-based cluster analysis to investigate the diversity of differentiation stage- and birthdate-specific cells and identified four embryonic age-defined AP transcriptional states, as well as two embryonic age-defined basal progenitor populations, as recently reported (Yuzwa et al. 2017) (Fig. 1C). Two classes of 1-day-old neurons (N1d) could be distinguished, early-born cells (*i.e.* E12-E13-born) and later-born cells (*i.e.* E14-E15-born). These two classes of neurons displayed early onset expression of deep- and superficial-layer markers, which foreshadowed their upcoming lamina-related identity (Fig. 1C). Classical deep-layer markers were also expressed by late-born neurons (fig S2A), consistent with an initial period of mixed identity followed by molecular cross-interactions and progressive fate refinement (Zahr et al. 2018; Ozair et al. 2018; Azim et al. 2009). Accordingly, by four days of age, neurons with mutually-exclusive expression of classical lamina-specific markers such as *Bcl11b* (an L5 marker), *Rorb* (L4) and *Cux1* (L2/3) emerged. Of note, GABAergic interneurons were also identified (Fig. 1C and fig. S2B), likely corresponding to cells migrating into the dorsal pallium after FT labeling of their progenitors in the ventral pallium (Govindan et al. 2018; Wamsley & Fishell 2017; Marin 2013). Astrocytes, corresponding to 4-day-old daughter cells of E15 APs (Minocha et al. 2017; Cahoy et al. 2008) were also present (Fig. 1C and fig. S2C). These two cell-types were not further investigated in this study. Differential expression analysis identified type-enriched transcripts (Fig. 1D) whose temporal patterns of expression were confirmed using *in situ* hybridization (Fig. 1E; figs. S3 and S4; ISH; Allen Developing Mouse Brain Atlas). Together, these results indicate that APs transit through temporally dynamic transcriptional states during corticogenesis as daughter neurons progressively acquire more mature transcriptional features.

We used two axes of investigation to address the transcriptional dynamics of APs and differentiating neurons: on the one hand, we studied the progression in AP transcriptional states between E12 and E15 (Fig. 2), and on the other hand we studied the transcriptional differentiation of neurons born on each of these embryonic days, as shown in Fig. 3. To address the temporal progression in AP transcriptional states we used a pseudotime (*i.e.* pseudo-birthdate) alignment approach (Mayer et al. 2018), which highlighted the chronotopic organization of these cells and identified clusters of genes with similar embryonic age-defined expression dynamics (Fig. 2, A and B; fig. S5, A and B). Several of these dynamically-expressed genes have previously characterized functions in the temporal regulation of progenitor competence, including the early-peaking transcripts *Hmga2* (E12 peak) and *Aspm* (E13 peak) and the late-peaking transcript *Zbtb20* (E15) (Kishi et al. 2012; Johnson et al. 2018; Tonchev et al. 2016). Ontological analysis revealed the progression of AP functional properties during corticogenesis (Fig. 2C, fig. S5C). At early embryonic ages (E12, E13), APs were involved in largely cell-autonomous tasks, such as regulation of gene expression and of chromatin structure; cell-death related processes were also prominent, suggesting some level of regulation of the size of the progenitor pool (Cunningham et al. 2013). Later in corticogenesis (E14, E15), external signaling and cell-cell interaction processes increased, as did lipid metabolism, which has been linked with progenitor fate in adult neuronal stem cells (Knobloch et al. 2013). Ion transport-related processes became more salient, in line with the role of bioelectrical parameters in the progression of AP competence (Vitali et al. 2018) and other typically neuron-related processes involving synapses and neurotransmission increased. As further discussed below, this suggests that APs progressively acquire molecular signatures of their progeny upon repetitive rounds of cell division. Finally, glia-related processes emerged, consistent with the generation of this cell type in late corticogenesis (Jabaudon 2017; Gao et al. 2014; Guo et al. 2013). Together, these findings identify the sequential unfolding of successive transcriptional and functional programs within APs as corticogenesis proceeds.

**Fig. 2.**
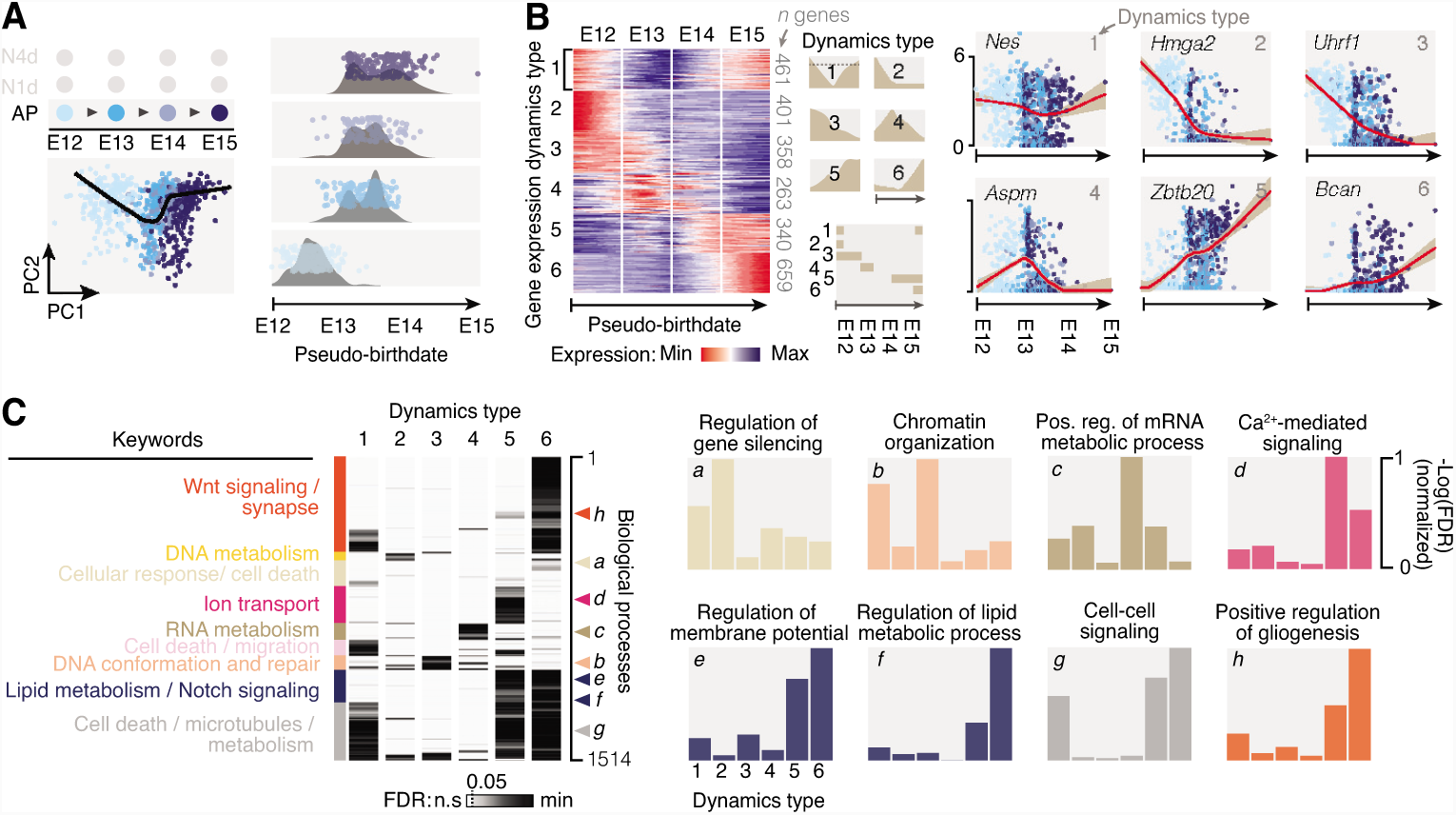
Apical progenitors transit through sequential transcriptional states during corticogenesis. (A) Principal component analysis (PCA) of AP transcriptional identity showing chronotopic organiza-tion along a birthdate axis (i.e. from E12 to E15). Cells were aligned on a pseudo-birthdate axis to trace this maturation route (black line). (B) Cluster analysis reveals distinct dynamics of AP gene expression during corticogenesis. Examples of genes for each type of dynamics are provided on the right. (C) Examples of gene ontology processes associated with each of the expression dynamics. Descriptions of functions in the left panel summarize relevant ontologies. Abbreviations: AP: apical progenitor, N1d: 1-day-old neuron, N4d: 4-day-old neuron, VZ: ventricular zone, CP: cortical plate.

**Fig. 3.**
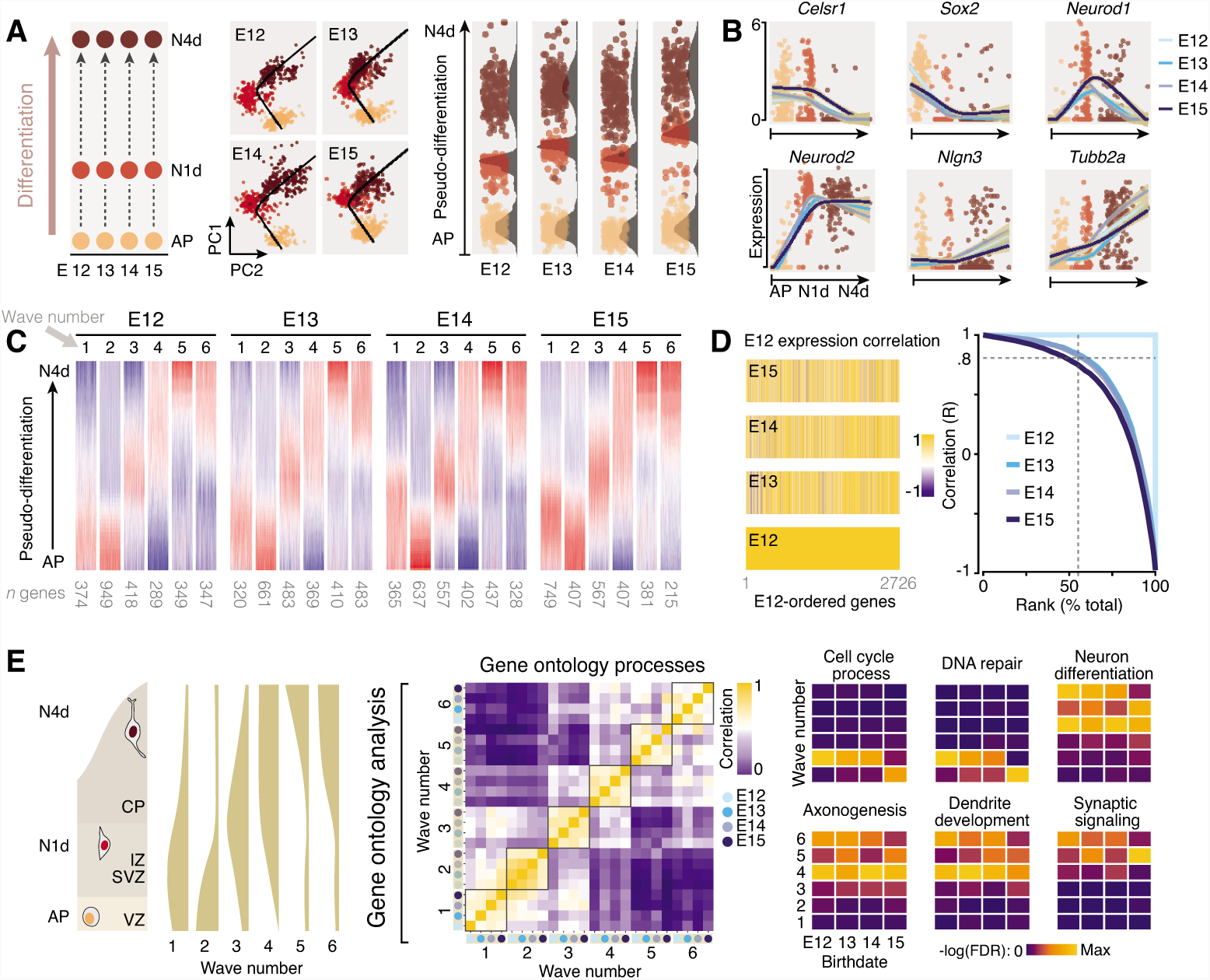
Neuronal differentiation programs are conserved across corticogenesis. (A) Principal component analysis (PCA) shows that at each developmental age, cells are spontaneously organized along a differentiation axis (i.e. from AP to N1d to N4d). Cells were aligned along a pseudo-differentiation axis to trace this maturation route (black line). (B) Gene expression kinetics along the pseudo-differentiation axis at each developmental age. Displayed cells correspond to E12. (C) Clustering of the gene expression kinetics reveals sequential expression of transcriptional waves. (D) Correlation of gene kinetics considering E12 as a reference shows that most gene expression dynamics are independent of the developmental age. (E) Parallel progression of gene ontology processes associated with each transcriptional wave. Abbreviations: AP: apical progenitor, CP: cortical plate, IZ: intermediate zone, N1d: 1-day-old neuron, N4d: 4-day-old neuron, SVZ: subventricular zone, VZ: ventricular zone.

We next examined the transcriptional programs expressed by differentiating neurons born on each embryonic age (Fig. 3). Pseudotime (*i.e.* pseudo-differentiation) alignment highlighted the sequential differentiation states of these cells and identified dynamically-expressed genes (Fig. 3 A-C; fig. S6, and see Methods). Clustering of these transcripts according to their expression dynamics outlined successive transcriptional waves driving differentiation (Telley et al. 2016) (Fig. 3C). The sequential unfolding of gene expression was essentially conserved across embryonic ages, as revealed by largely overlapping gene expression dynamics (Fig. 3D and fig. S6). Conserved gene expression did not simply reflect the constant activity of a small number of “pan-neuronal” genes (*e.g.* NeuroD2, Tubb2) but instead reflected genuinely conserved differentiation programs since over half of the expressed genes had highly conserved expression dynamics (R > 0.8, Fig. 3D). Accordingly, gene ontologies within waves were conserved across embryonic ages (Fig. 3E). Thus, in contrast to the programs driving the temporal progression in AP identity, the differentiation programs of daughter neurons are largely conserved across embryonic ages, despite the distinct identities these daughter neurons acquire.

How then does neuronal diversity emerge? As reported above, the chronotopic arrangement of APs is also present in their 1-day old progeny (Fig. 1B). This suggests that embryonic age-dependent AP transcriptional programs are transmitted to their progeny to generate successive initial neuronal identities. To investigate this possibility, we next determined how dynamic transcriptional networks emerge in single cells during corticogenesis. We used a machine learning strategy to classify cells based on (1) their birthdate and (2) their differentiation status, which identified two core sets of genes (*n* = 100 per model) sufficient to classify all cells according to these two cardinal features (Fig. 4A and fig. S7A-C), many of which have been previously involved in regulating progenitor and neuronal fate (Tables 1 and 2). Birthdate-associated core genes were sequentially expressed by APs and their 1-and 4-day old progeny, directly demonstrating transmission of age-specific genesets to daughter cells (Fig. 4B, top and fig. S7D). In contrast, consistent with a consensus post-mitotic differentiation program, the dynamics of the differentiation geneset were conserved across embryonic ages (Fig. 4B, bottom). Consistent with the increase in neuron-related ontologies in APs noted above, expression of the neuronal differentiation geneset progressively increased in APs as corticogenesis unfolded (fig. S7E and F); the latter cells thus become progressively “neuralized” as they give rise to successive generations of post-mitotic daughter cells. Taken together, these data reveal that neuron-type-specific identities emerge from temporally-defined, AP-derived transcriptional ground states onto which essentially conserved post-mitotic differentiation programs are applied (Fig. 4C).

**Table 1:**
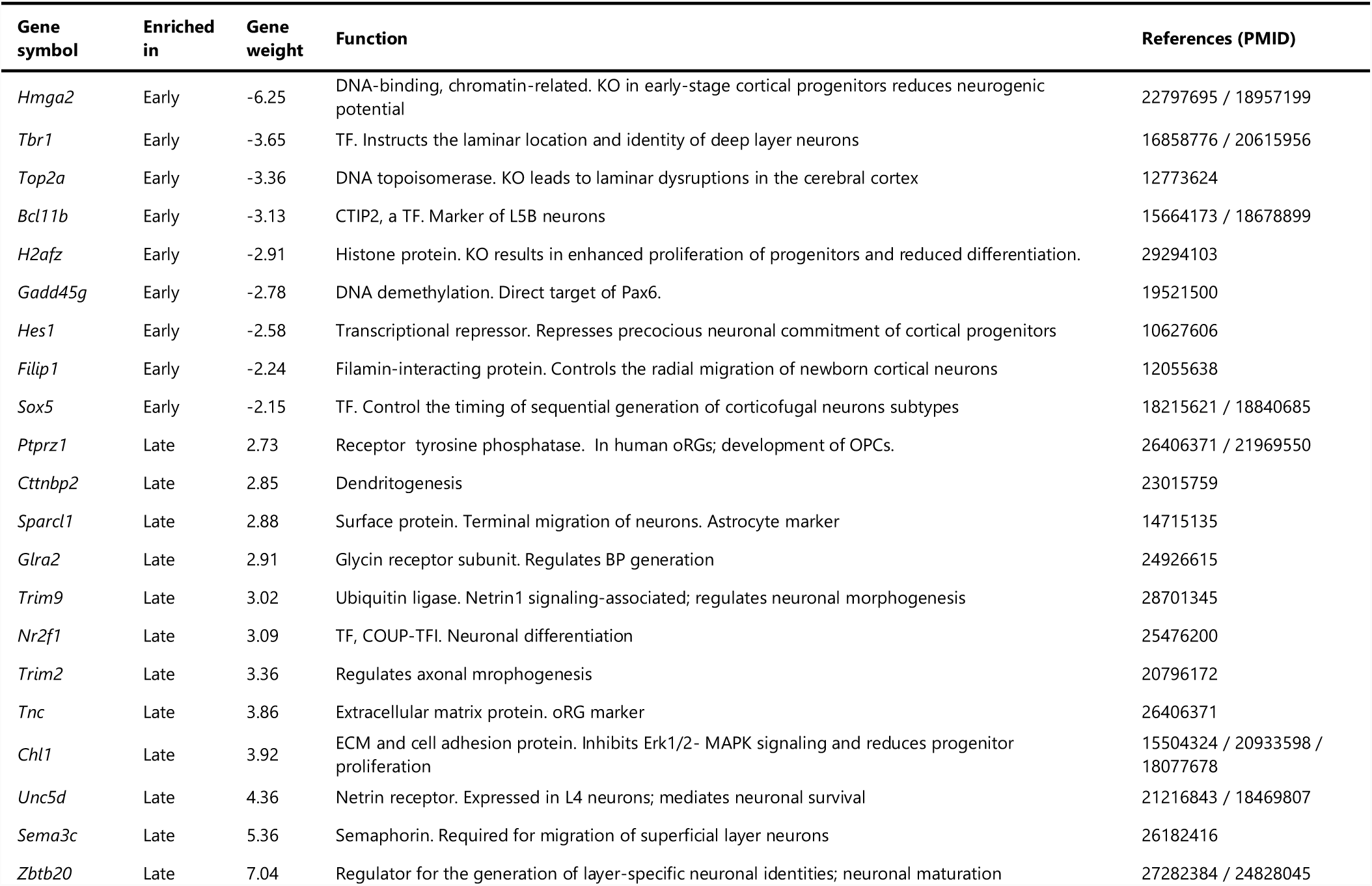
Selection of characterized genes from the birthdate model

**Table 2:**
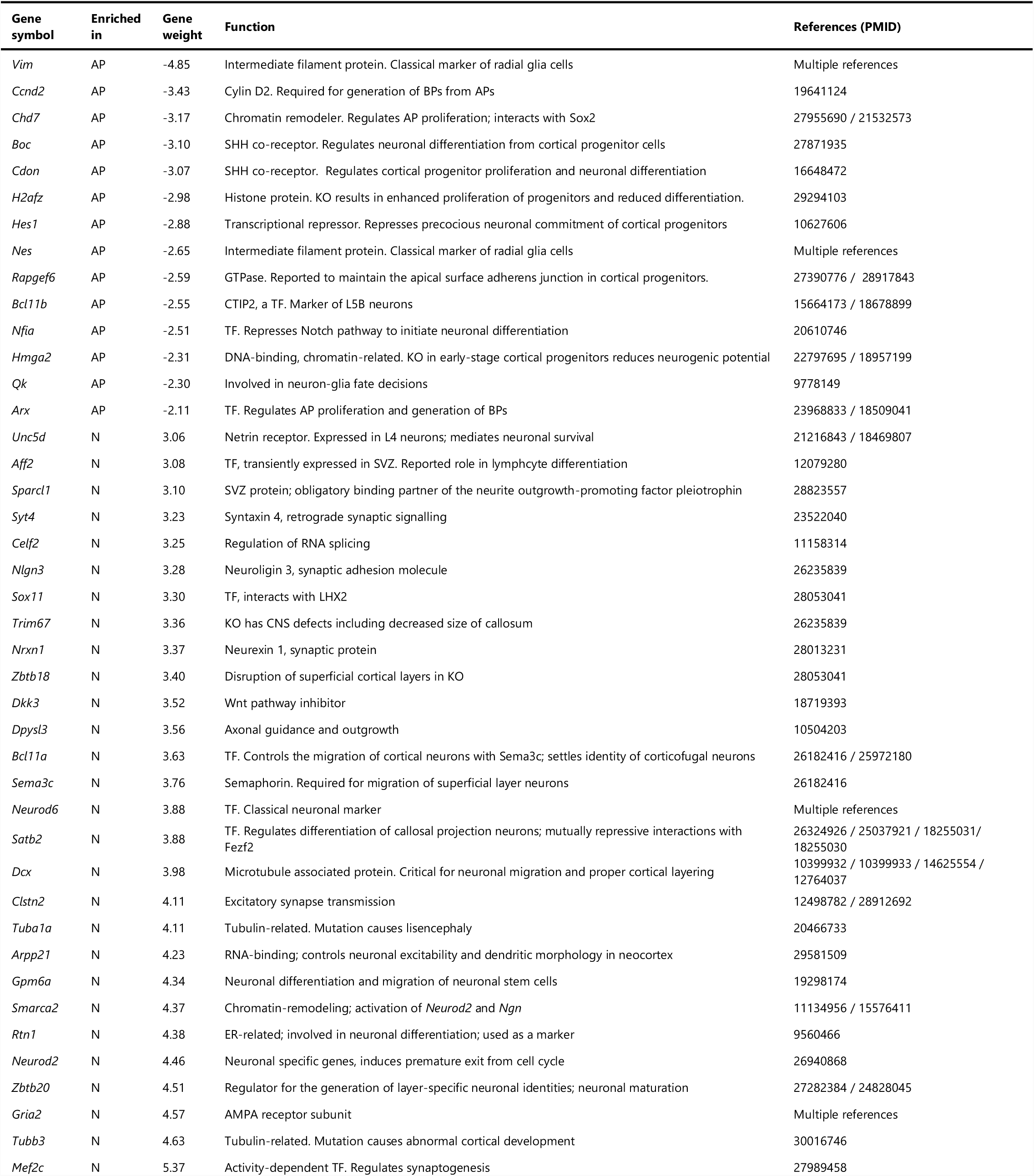
Selection of characterized genes from the differentiation model

**Fig. 4.**
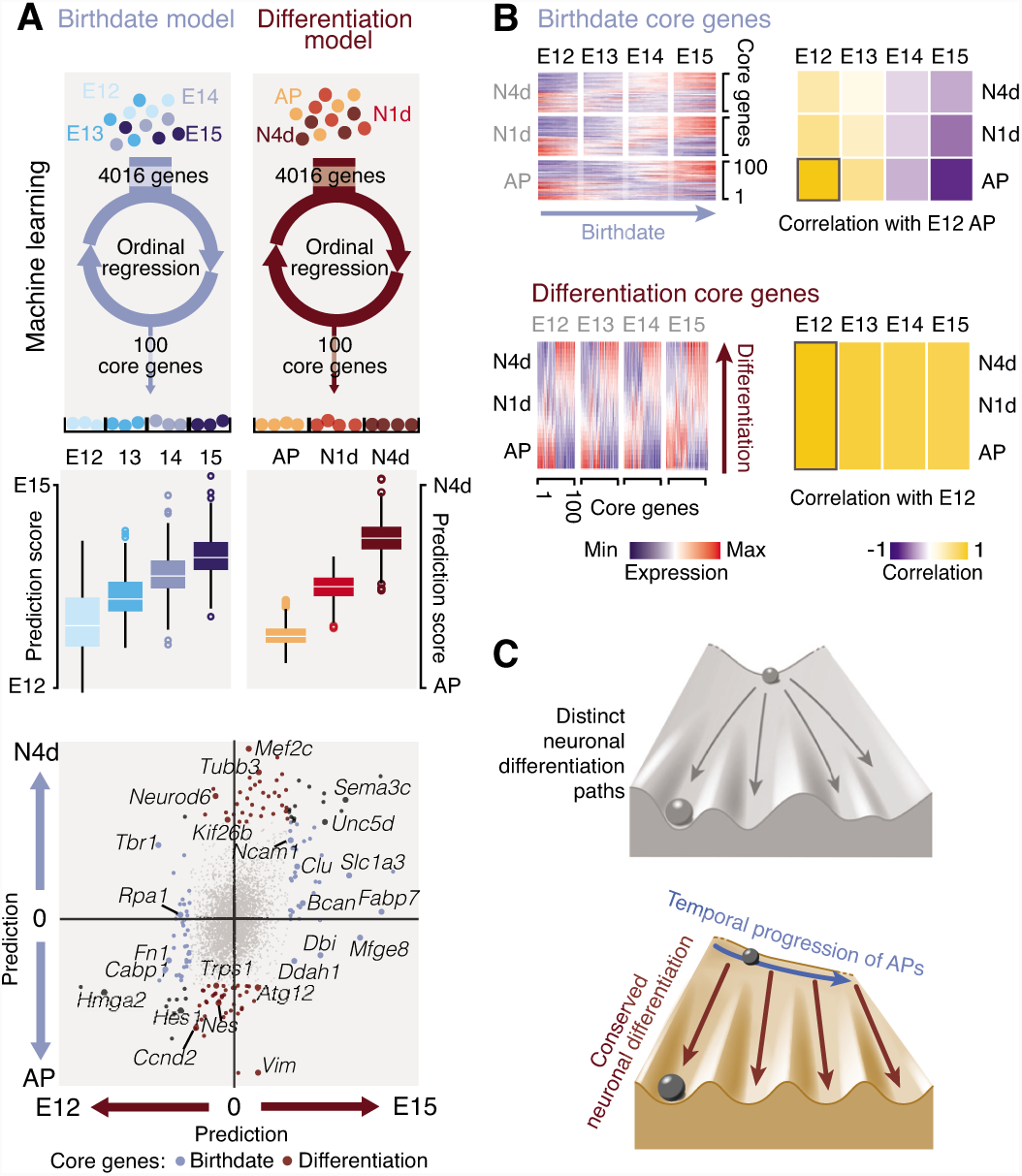
Temporally progressing AP transcriptional states interact with conserved differentiation programs to generate neuronal diversity. (A) Machine learning approach used to identify a core set of genes which can classify cells based on their date of birth (top left) and differentiation status (top right). Center: Model performance using actual dataset. Box plot: median ± SEM. Bottom: Weight of the core genes in predicting birthdate and differentiation status. See also tables S1 and S2. (B) Top: Birthdate-associated core genes are temporally dynamic and daughter cells acquire embryonic stage-specific transcriptional birthmarks. See also Supplementary Fig.7D. Bottom: In contrast, differ-entiation status-associated core genes are conserved across corticogenesis. (C) Schematic representation of the findings: in the classical Waddington epigenetic model (top) cellular diversity emerges through distinct developmental trajectories. The current data shows that instead, in the neocortex, developmental trajectories are conserved, but that initial ground states are temporally dynamic. Abbreviations: AP: apical progenitor, N1d: 1-day-old neuron, N4d: 4-day-old neuron.

We combined the two aforementioned models to identify birthdate- and differentiation stage-related patterns of gene expression. Based on the combined expression of the core genes of the two models, each cell was assigned a birthdate score and differentiation score. Cells were then embedded within a two-dimensional matrix, allowing the display of gene expression profiles as chrono-typic transcriptional maps (Fig. 5A) (Nowakowski et al. 2017). This approach revealed a variety of dynamically-regulated transcriptional patterns, including within single families of genes (Fig. 5B and fig. S8). To identify archetypical features of gene expression, we performed a t-SNE-based cluster analysis of all transcriptional maps, revealing canonical clusters of genes with similar expression dynamics (Fig. 5C). Genes within each of these canonical clusters shared common functions, and the distinct clusters were functionally specialized (Fig. 5D and fig. S9). This suggests that these transcriptional clusters represent functional units orchestrating the unfolding of cellular processes during corticogenesis. To substantiate this possibility, we selected one early and one late AP transcriptional process and assessed its functional outcome (Fig. 5E and F). Expression of the Polycomb Repressive Complex 2 (PRC2), which regulates histone methylation in neural progenitors (Pereira et al., 2010), provided a first example: all three subunits of the complex were co-expressed in APs early in corticogenesis, and the H3K27me3 signature mark of PRC2 had corresponding dynamics on target sites, demonstrating temporally-gated functional activity (Fig. 5E). Expression of the glutamate transporter transcript *Slc1a3* (*Glast*) constituted a second example: *Glast* increased in APs as glutamatergic neurotransmission developed in the cortical plate. Pharmacological blockade of this transporter increased glutamate levels at late, but not early embryonic stages, consistent with a dynamic bioelectrical control over AP properties during corticogenesis (Fig. 5F) (Vitali et al. 2018).

**Fig. 5.**
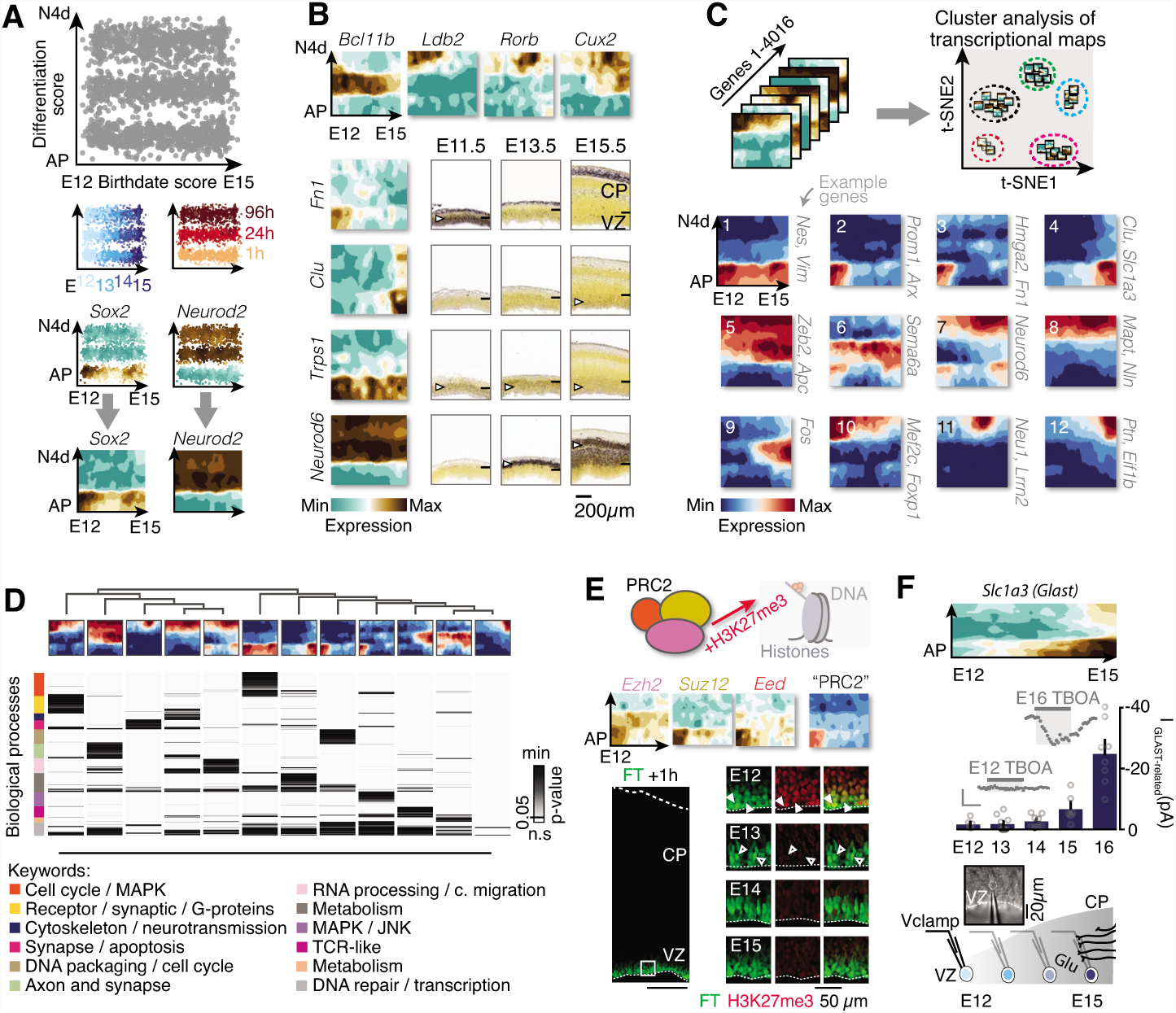
Dynamic transcriptional mapping of corticogenesis. (A) Cellular map in which cells are displayed based on their combined expression of the birthdate and differentiation status core genes presented in Fig. 4. Dynamic expression of genes (“transcriptional maps”) throughout corticogenesis can be determined based on this template (as exemplified with Sox2 and NeuroD2). (B) Example of transcriptional landscapes for select genes, and corresponding ISH validation from the Allen Developing Mouse Brain Atlas (www.brain-map.org). (C) Canonical transcriptional maps can be identified by t-SNE clustering of the maps of individual genes. (D) Genes belonging to each of the clusters have converging and specialized ontologies. (E) The distinct components of the polycomb repressive complex 2 have equivalent chrono-typic expression profiles; functional interrogation of this complex with the H3K27me3 mark reveals a corresponding methylation enrichment in E12 APs. (F) The glutamate transporter transcript Slc1a3 (Glast) is expressed by APs late in corticogenesis. Blockade of GLAST with TBOA (Jabaudon et al. 1999) increases extracellular glutamate at late but not early embryonic ages, as detected by activation of ionotropic glutamate receptors in patch-clamped APs. Scale bars: 5 min; 20 pA. Abbreviations: AP: apical progenitor, N4d: 4-day-old neuron, PRC2: Polycomb Repressive Complex 2, CP: cortical plate, VZ: ventricular zone.

Together, our findings identify a combinatorial process in which type-specific neuronal identity emerges from the apposition of generic differentiation programs onto embryonic age-dependent, AP-derived transcriptional states. In this scenario, neuronal differentiation essentially corresponds to the implementation of programs coding for generic neuronal features (*e.g.* neurites, neurotransmission) onto temporally-defined initial transcriptional states. This process is reminiscent of how neuron diversity is generated in evolutionary older brain regions such as the subpallium or spinal cord (Mayer et al. 2018; Mi et al. 2018; Nowakowski et al. 2017; Dasen & Jessell 2009), with the difference that in these regions, distinctions in initial neuronal states reflect a predominantly spatial rather than temporal distribution of molecularly distinct progenitors. There thus appears to be at least two ways to generate cellular diversity: spatial patterning of molecularly distinct progenitors (*e.g.* subpallium, spinal cord), and temporal patterning, as revealed here. In evolutionary terms, temporal patterning may have been selected as the primary mode of neuron production in the neocortex because it allows the generation of a large spectrum of cell types at low spatial cost. How temporal birthmarks are transmitted from APs to daughter neurons is unclear, but the strong temporal control over epigenetic processes identified here suggests that transmission of 3D chromatin features may be involved. In addition, the passive transmission of cytoplasmic RNA into daughter neurons along with post-transcriptional events could also contribute (Zahr et al. 2018; Yoon et al. 2017, 2018). As previously reported, we find that newborn neurons initially express a combination of classical lamina type-specific markers, a process which has been termed “transcriptional priming” and is also found in other organs such as the hematopoietic system (Zahr et al. 2018; Azim et al. 2009; Hu et al. 1997). Our findings thus do not exclude the contribution of post-mitotic processes to fate refinement (Zahr et al. 2018; Ozair et al. 2018; Mayer et al. 2018; Telley & Jabaudon 2018), but AP-derived, temporally-regulated processes appear to have a primordial role in defining initial neuronal identity. Although still present in 4-day-old neurons, temporal birthmarks fade with differentiation. At these later stages, activity-dependent programs may be progressively implemented in interaction with the environment to complement and eventually override earlier transcriptional processes, culminating in the generation of the full complement of cells required for functional cortical circuits.

## Acknowledgments

We thank A. Benoit and the Genomics Platform and FACS Facility of the University of Geneva for technical assistance; and H. Wu, E. Azim, O. Raineteau, D. Silver and the members of the Jabaudon laboratory for comments on the manuscript. Illustration in panel 4C: www.lagraphisterie.fr. Work in the Jabaudon laboratory is supported by the Swiss National Science Foundation and the Carigest Foundation. DJ and AD are supported by the Fondation privée des HUG. G.A. is a PhD student from the F.R.S-F.N.R.S and is supported by the Fonds Léon Fredericq. The LN laboratory is funded by the F.R.S.-FNRS (EOS O019118F-RG36; CDR J.0028.18; PDR T.0073.15) the Fonds Léon Fredericq, the Fondation Médicale Reine Elisabeth, and the Fondation Simone et Pierre Clerdent. A.D. is funded by the Swiss National Science Foundation and the National Center of Competence in Research (NCCR) Synapsy.

## Author contributions

LT and GA performed the experiments with the help of PO and IV. SF performed the electrophysiology experiments. LT, GA and JP performed the bioinformatic analysis. GA, DJ and LT wrote the manuscript with the help of AD and LN.

## Data and materials availability

Annotated data will be available upon manuscript publication. The supplementary materials contain additional data.

## Materials and Methods

### Mice

All experiments were approved by the Geneva Cantonal Veterinary Authorities, Switzerland. To avoid developmental variability between embryos, three hour-time-mated pregnant CD1 mice were purchased from Charles River Laboratories. Embryonic day (E) 0 was established as the time of detection of the vaginal plug. Both female and male embryos were analyzed in this study.

### *In utero* FlashTag injection

FlashTag (FT) injections were performed at E12, E13, E14 or E15, as previously described (Telley et al. 2016; Govindan et al. 2018). Briefly, pregnant females were anaesthetized with isoflurane, treated with Temgesic (Reckitt Benckiser, Switzerland) and the uterine horn was exposed following an abdominal incision. Half a microliter of 10 mM of a carboxyfluorescein succinimidyl ester (*i.e.* Flash-Tag, CellTrace™ CFSE, Life Technologies, #C34554) was injected into the lateral cerebral ventricle of the embryos. The abdominal wall was then closed and the embryos were let to develop until collection.

### Immunofluorescence and imaging

#### Tissue processing

Embryonic brains were dissected in a phosphate-buffered saline (PBS) solution, fixed in 4% paraformaldehyde (PFA) overnight at 4 °C then cryoprotected in PBS-sucrose 30% overnight at 4°C before embedding in OCT and freezing on dry ice. On-slide coronal brain sections with a thickness of 14 μm were performed using a cryostat.

#### Immunofluorescence on brain sections

Brain sections were post-fixed 10 min in 4% PFA, washed three times in PBS, incubated 30min at 85 °C in citrate buffer solution and washed 3 times in PBS prior to a 1-hour incubation in blocking solution (10% horse serum - 0,5% Triton X-100 diluted in PBS) at room temperature. Slides were then incubated overnight at 4°C with primary antibodies. Next, slides were washed 3 times in PBS and incubated 2 hours at room temperature with respective secondary antibodies (1:500) before mounting with Fluoromount (Sigma). Primary antibodies used: rabbit anti-pH3 (1:500, Abcam, #AB5176), rabbit anti-H3K27me3 (1:500, Millipore, 07-449).

### Imaging

All images were acquired on LSM 700 confocal laser scanning microscope (Carl Zeiss). The putative primary somatosensory (S1) cortex was used as a region of study. The ImageJ software was further used for downstream image processing.

### *In situ* hybridization image processing

All *in situ* hybridizations were retrieved from the Allen Developing Mouse Brain Atlas (www.brain-map.org) and uniformly zoomed to the putative S1 neocortical region. For the illustrations Fig. 1E and figs S4, S5C, S7C the images were aligned and stacked. The mean intensity level of the Z projection was calculated on ImageJ. The resulting layout was artificially colored using the “Fire” mode of ImageJ.

### scRNAseq experiment

#### Cell dissociation and FAC-sorting

Pregnant females were sacrificed either 1, 24 or 96 hours after FT injection. As previously described (Telley et al. 2016; Govindan et al. 2018), embryonic brains were extracted in ice-cold HBSS, embedded in 4% agar low-melt and sectioned coronally at 300 μm using a vibrating microtome (Leica, #VT100S). The putative S1 cortical region was microdissected under a stereomicroscope and incubated in 0.05% trypsin at 37°C for 5 minutes. Following tissue digestion, fetal bovine serum was added to the mix and cells were manually dissociated via up-and-down pipetting. Cells were centrifuged 5 min at 300 G and the pellet was suspended in 1 ml of HBSS then passed on a 70 μm cell strainer. FT^+^ cells, gated to include only the top 5% brightest cells (Telley et al. 2016; Govindan et al. 2018), were finally FAC-sorted on a MoFloAstrios device (Beckman).

#### Single-cell RNA capture and sequencing

FAC-sorted FT^+^ cells (18 μl) were mixed with the C1 Suspension Reagent (2 μl; Fluidigm) yielding a total of 20 μl of cell suspension mix with ∼500 cells / μl. The cell suspension mix was loaded on a C1 Single-Cell AutoPrep integrated fluidic circuit (IFC) designed for 10- to 17-μm cells (HT-800, Fluidigm #100-57-80). cDNA synthesis and preamplification was processed following the manufacturer’s instructions (C1 system, Fluidigm) and captured cells were imaged using the ImageXpress^®^ Micro Widefield High Content Screening System (Molecular Devices^®^). Single cell RNA-sequencing libraries of the cDNA were prepared using Nextera XT DNA library prep kit (Illumina). Libraries were multi-plexed and sequenced according to the manufacturer’s recommendations with paired-end reads using HiSeq2500 plat-form (Illumina) with an expected depth of 1M reads per single cell, and a final mapping read length of 70 bp. All the single cell RNA capture and sequencing experiments were performed within the Genomics Core Facility of the University of Geneva. The sequenced reads were aligned to the mouse genome (GRCm38) using the read-mapping algorithm TopHat. Unique Molecular Identifiers (UMI) sequenced in the first reads were used to correct for cDNA PCR amplification biases. Duplicated reads were identified and corrected using the deduplication step from the UMI-tools algorithm (doi:10.1101/gr.209601.116). The number of reads per transcript was calculated with the open-source HTSeq Python library. All the analyses were computed on the Vital-It cluster administered by the Swiss Institute of Bioinformatics.

### scRNAseq analysis

#### Cell filtering

Doublet cells identified on the Fluidigm C1 plate images were excluded before initial analysis. A total of 2,906 FT^+^ single cells were obtained (**FT +1 h**: E12: 202 cells, E13: 211, E14: 135, E15: 304; **FT +24 h:** E12: 284 cells, E13: 286, E14: 232, E15: 217; **FT + 96 h:** E12: 246 cells, E13: 278, E14: 262, E15: 249). Cells expressing < 1000 genes or > 17% of mitochondrial genes were excluded. After this step, 2’756 cells remained for analysis (**FT +1 h**: E12: 189 cells, E13: 207, E14: 134, E15: 301; **FT +24 h:** E12: 268 cells, E13: 223, E14: 219, E15: 213; **FT +96 h**: E12: 244 cells, E13: 267, E14: 254, E15: 237).

#### Type specific transcripts

The AP, BP and N score used in Fig. 1B correspond to the mean transcript expression of the top 20 genes for AP, BP and N previously characterized in (Telley et al. 2016) were: **AP**: *Aldoc, Pdpn, Vim, Ednrb, Ddah1, Ldha, Peg12, Wwtr1, Tspan12, Mfge8, Uhrf, Ncaph, Ndrg2, Mt1, Hk2, Psat1, Sp8, Sdc4, Dnmt3a, Notch2, Psph*. **BP**: *Btg2, Eomes, Abcg1, Kif26b, Mfap4, Coro1c, Myo10, Mfng, Rprm, Chd7, Ezr, Gadd45g, Slc16a2, Heg1, Celsr1, Tead2, Cd63, Rhbdl3, Mdga1, Arrdc3*. **N**: *Myt1l, Unc5d, 1700080N15Rik, Nos1, Satb2, Ank3, Scn3a, Dscam, Cntn2, Plxna4, 9130024F11Rik, Lrrtm4, Ptprk, Nrp1, Celsr3, Rbfox1, Flrt2, Kcnq3, Kcnq2, Gm36988.*

*Clustering analysis* was performed using the Seurat bioinformatics pipeline (https://github.com/satijalab/seurat) and is summarized here. We first created a “Seurat object” including all 2,756 cells and all genes. To remove sequencing depth biases between cells, we normalized and scaled the UMI counts using the *NormalizeData* (normalization.method = “LogNormalize”, scale.factor = 100000) combined with the *ScaleData* function (vars.to.regress = c(“nGene”,”nUMI”)). We then determined the most variable genes by plotting transcripts into bins based on X-axis (average expression) and Y-axis (dispersion). This identified 4,016 transcripts. Parameters and cutoffs were set as follow: mean.function = ExpMean, dispersion.function = LogVMR, x.low.cutoff = 0.1, x.high.cutoff = 8, y.cutoff = 0.7. Next, we identified the statistically significant principal components and used the top 20 as input for t-Distributed Stochastic Neighbor Embedding dimensional reduction, using the *TSNEPlot* function. To identify cellular clusters, we adopted a graph-based clustering approach using *FindClusters* function with a 1.8 resolution. Finally, a multiclass SVM model (implementation from R package *bmrm* was trained on all cluster-assigned cells and genes were ranked according to their linear weights. For each class (*i.e.* clusters), genes with a significant linear weight (Z-test, FDR ≤ 0.05) were considered as enriched genes.

#### Pseudotime projection

APs, N1d and N4d cells at all embryonic ages identified in the cell clustering analysis were processed. Basal progenitors were not included in this analysis because N1d and N4d are overwhelmingly directly born from APs when using FT labeling (Telley et al. 2016; Govindan et al. 2018). The pseudotime alignment method performed was previously described (Mayer et al. 2018) and is summarized hereafter. In Fig. 2, Fig. 3 and figs. S5 and S6, we restricted the datasets to the high variable genes (*n* = 4,016) and performed dimensionality reduction using the prcomp function of R software. Taking into consideration the significant principal components (PCs) explaining at least 2% of the total variance and using the R package princurve, we fitted a curve that described the maturation route (*i.e.* pseudo-birthdate or pseudo-differentiation) along which cells are organized. The beginning of the curve was established as the side where cell expressed the highest level of *Sox2* (AP) for pseudo-differentiation or the highest level of *Hmga2* (E12) for pseudo-birthdate. A maturation score reflecting the distance beginning of the curve-cell coordinate was attributed to each cell and normalized between 0 to 1. We then restricted the dataset to the top 500 genes for each PCs and performed a “Partitioning Around Medoids” analysis using the PAM R package (K = 6, span = 0.6) to identify clusters of transcripts with similar expression dynamics along the pseudo-birthdate (Fig. 2, fig. S5) or pseudo-differentiation (Fig. 3, fig. S6). This approach was previously described elsewhere (Telley et al. 2016).

#### Ordinal regression models

We used an ordinal regression method to predict on one hand the birthdate and on the other hand the differentiation status of each cell. We restricted the analysis to the high variable genes (*n* = 4,016) defined earlier. As the cells are expected to be organized within a differentiation and a birthdate continuum, we used and adapted a previously described ordinal regression model (Teo et al. 2010) implemented in the *bmrm* R package. In our context, a single linear model is optimized to predict cell differentiation status irrespectively of the date of birth and conversely. The linear weight of the models is used to rank the genes according to their ability to predict each cell category and the best 100 genes in each model were considered. The ordinal regression models were then re-optimized on these subsets of genes. All reported predictions were obtained by 10-fold cross-validation.

#### Transcriptional maps

(Fig. 5): Cells were organized on a 2D grid based on their birthdate and differentiation status score. For this purpose, the data were linearly adjusted so that the average predicted values for each cardinal feature was aligned on to the relative knot of the grid. The gene expression at a given coordinate of the 2D space was further estimated as the average expression of its 15 nearest neighbors. All transcriptional landscapes of the most variable genes (*n* = 4,016) were further clustered by projecting genes onto a 2D t-SNE space and submitted to a k-means clustering (K = 12).

### Electrophysiology

Four hundred μm-thick coronal slices were prepared from E12.5, E13.5, E14.5, E15.5 and E16.5 CD1 mice embryos and kept 30 minutes at 33°C in artificial cerebrospinal fluid (aCSF) containing 125 mM NaCl, 2.5 mM KCl, 1 mM MgCl_2_, 2.5 mM CaCl_2_, 1.25 mM NaH_2_PO_4_, 26 mM NaHCO_3_ and 11 mM glucose, saturated with 95% O_2_ and 5% CO_2_. Slices were then transferred in the recording chamber, submerged and continuously perfused with aCSF. The internal solution used for the experiments contained 140 mM potassium methansulfonate, 2 mM MgCl_2_, 4 mM NaCl 0.2 mM EGTA, 10 mM HEPES, 3 mM Na_2_ATP, 0.33 mM GTP and 5 mM creatine phosphate (pH 7.2, 295 mOsm). Cells in immediate proximity to the ventricular wall (i.e. putative APs) were patched and clamped at -70mV. A baseline stable holding current was first measured for 4 minutes, after which a 10-minute bath of 100 μM of the glutamate transporter antagonist _DL_-TBOA (DL-*threo*-β -Benzyloxyaspartate) (Jabaudon et al. 1999) was applied and finally washed out. TBOA-induced currents were blocked by application of 25 μM NBQX and 50 μM D-APV (data not shown), consistent with activation of ionotropic glutamate receptors by increased extracellular levels of Glu (Jabaudon et al. 1999). Recorded currents were amplified (Multiclamp 700, Axon Instruments), filtered at 5kHz, digitalized at isochronic cohorts of cells in20kHz (National Instrument Board PCI-MIO-16E4, IGOR WaveMetrics), and stored on a personal computer for further analyses (IGOR PRO WaveMetrics). The net amplitude of TBOA induced currents was determined after subtraction of baseline holding current. Values are represented as mean ± SEM.

### Softwares

All single cell RNA sequencing analysis were perfomed using the R software with publicly available packages. GeneGo portal (https://portal.genego.com) was used to investigate the enriched gene ontology processes in Fig. 2 and Fig. 3 and the biomart R package served to extract the list of genes allocated to a defined ontology term. Cytoscape platform (Maere et al. 2005) associated with its plugin (Shannon et al. 2003) was used to construct the enrichment gene ontology processes network in supplementary fig. S9. For this purpose, the latest version of gene ontology (go-basic.obo) and gene association (gene_association.mgi) from the Gene Ontology Consortium website (www.geneontology.org) were given as input in Bingo. The string database (http://string-db.org) implemented in Cytoscape platform was used to determine the protein-protein interactions in figs S5, S6 and S7.

## Supplementary figures

**Fig. S1.**
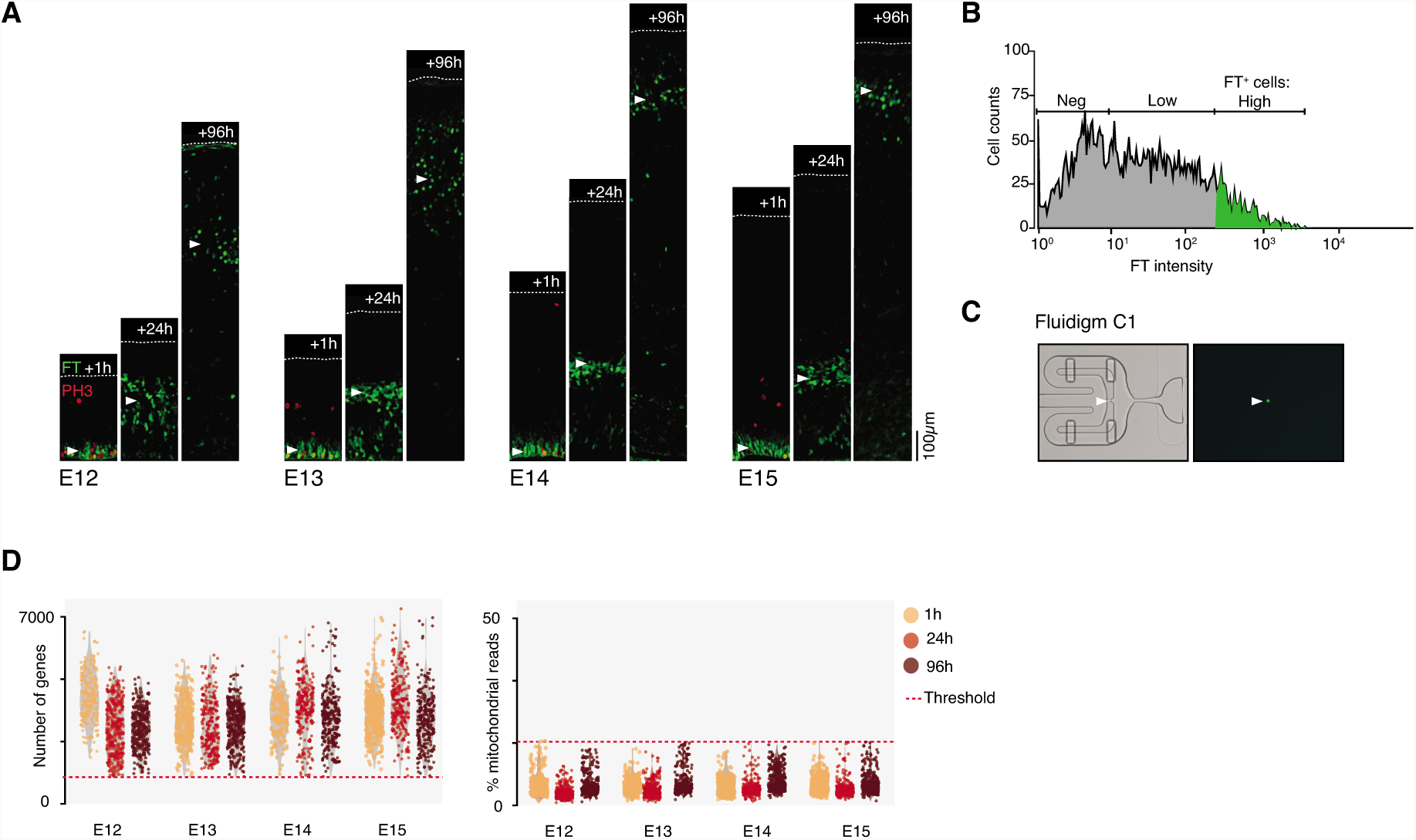
scRNA sequencing of isochronic cohorts of cells in the developing neocortex. (A) FT injection labels isochronic cohorts of cortical cells which were collected either 1, 24 or 96 hours after labeling. E12 illustration modified from Fig.1. (B) FT+ cells were FAC-sorted and (C) captured on a C1 microfluidic device for scRNAseq processing. (D) Violin plots showing the number of genes and the percentage of mitochondrial genes detected in each single cell. Lower (< 1000 genes) and upper (> 17% mitochondrial genes) accepted thresholds are displayed.

**Fig. S2.**
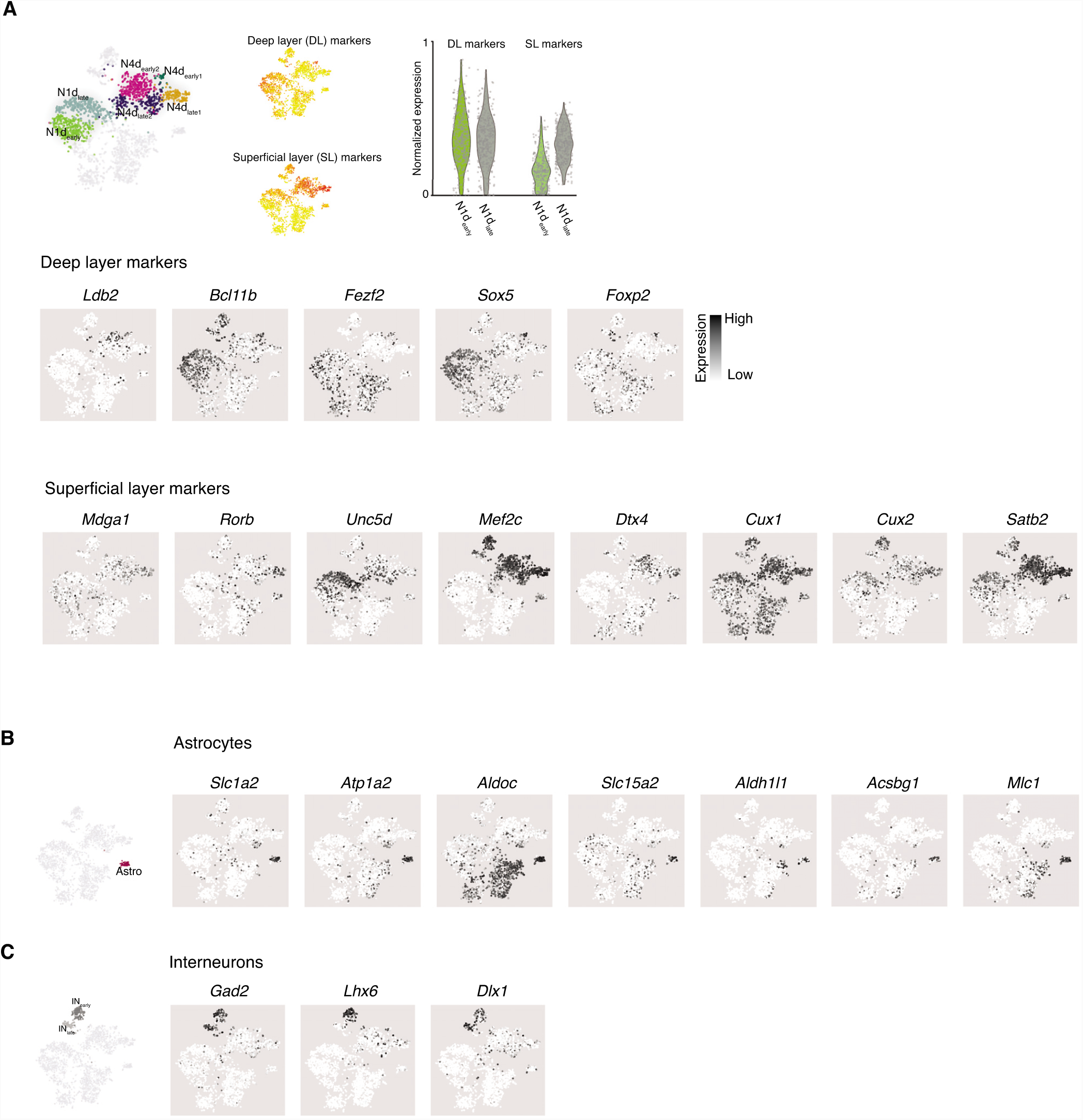
Expression of select genes in neurons, astrocytes, and interneurons. (A) Feature plots showing expression of classical deep layer (DL) and superficial layer (SL) markers in 1-day and 4-day-old neurons across corticogenesis. Note that N1dlate neurons (i.e. E14 and E15-born neurons) initially and transiently express DL markers. (C) Expression of select astrocytes markers. (D) Expression of select ventral pallial-derived interneuron markers.

**Fig. S3.**
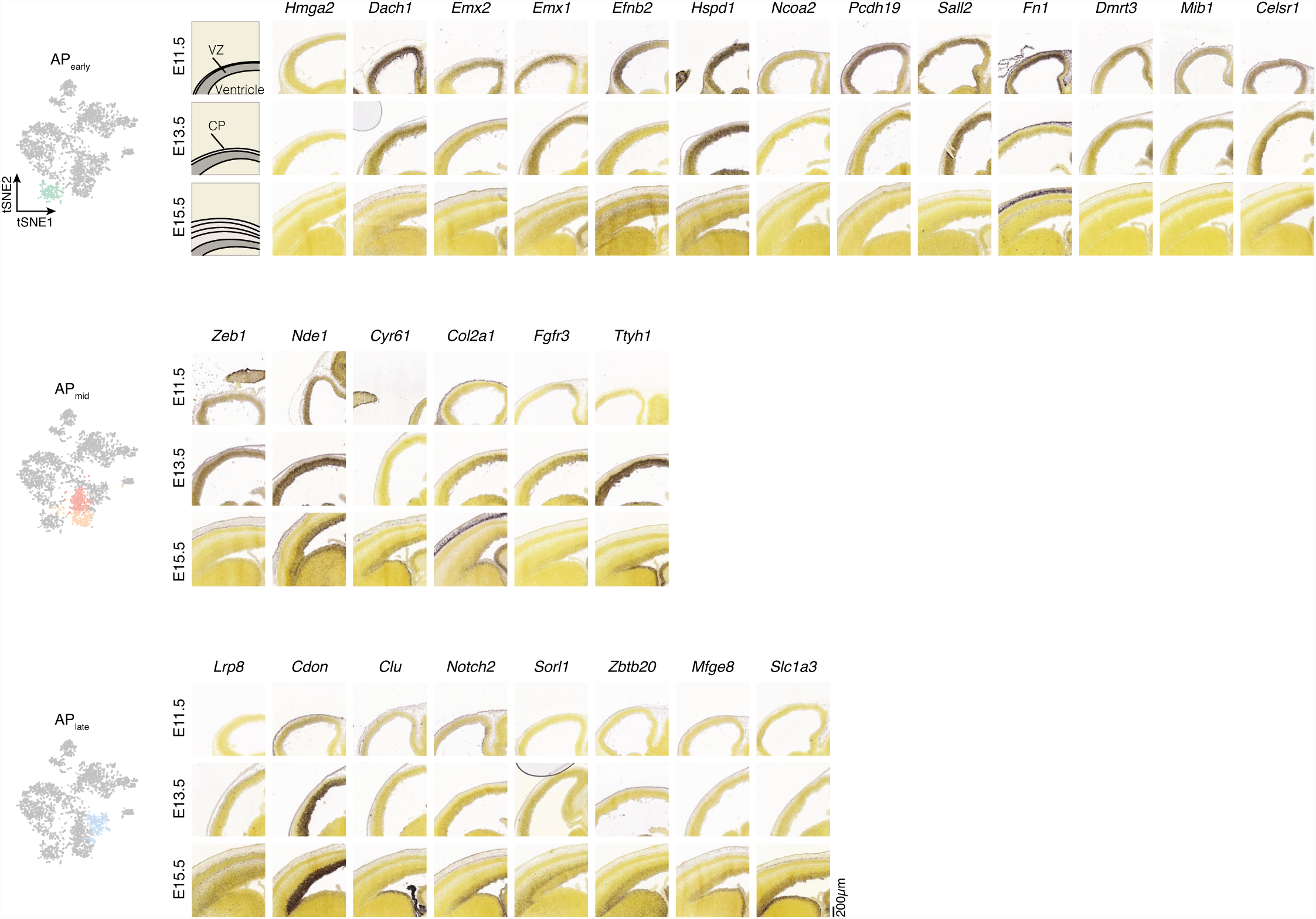
Spatio-temporal expression of the most enriched genes in AP clusters. In situ hybridization of genes enriched in APearly, APmid, and APlate clusters. The ISH merged layouts are also presented in Fig. 1E. Source of ISH: Allen Developing Mouse Brain Atlas.

**Fig. S4.**
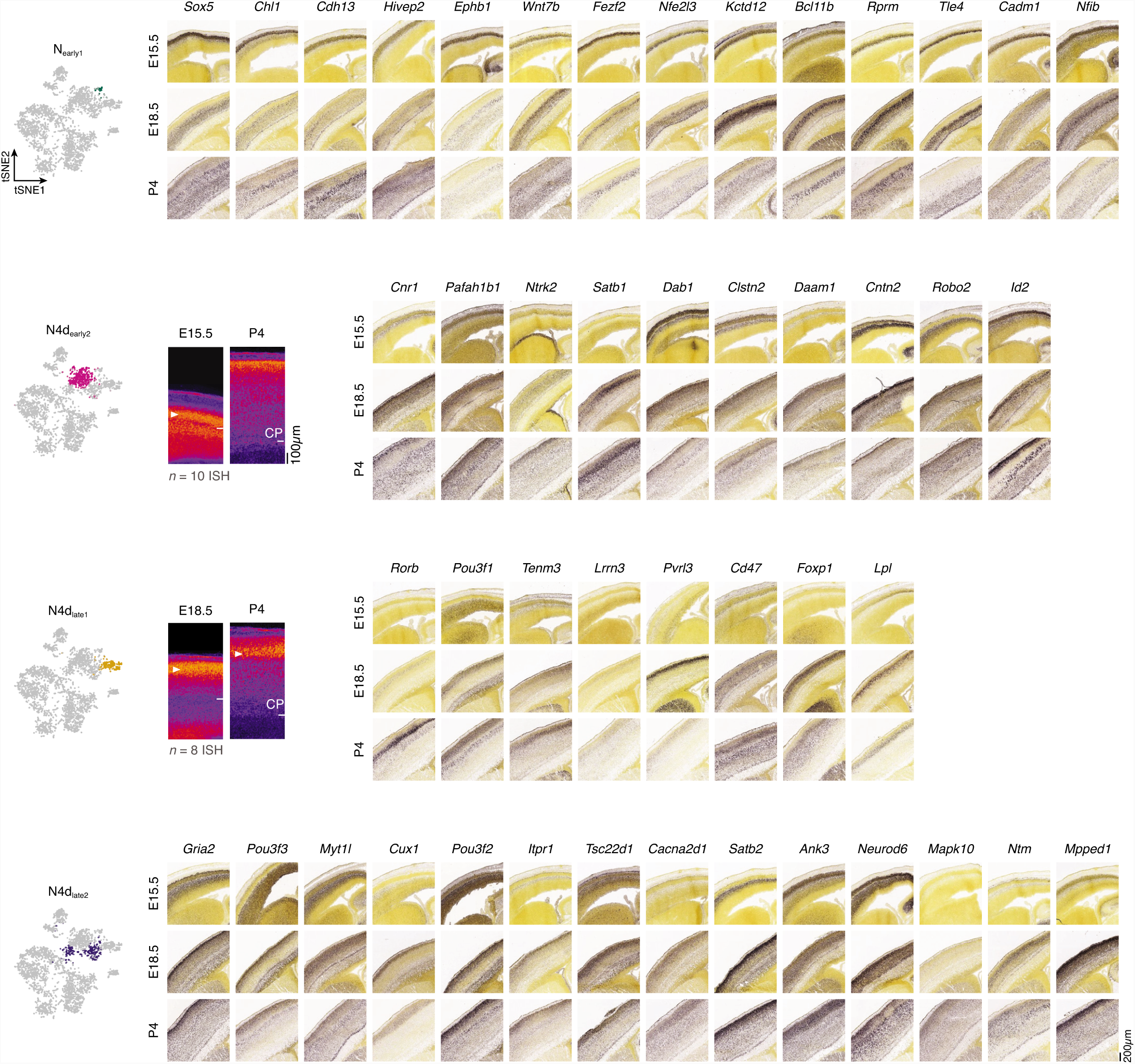
Spatio-temporal expression of the most enriched genes in 4-day-old clusters. Individual and merged *in situ* hybridization of genes enriched in N4dearly1, N4dearly2, N4dlate1 and N4dlate2 clusters. The ISH merged layouts for N4dearly1 and N4dlate2 are also presented in Fig. 1E. Source of ISH: Allen Developing Mouse Brain Atlas.

**Fig. S5.**
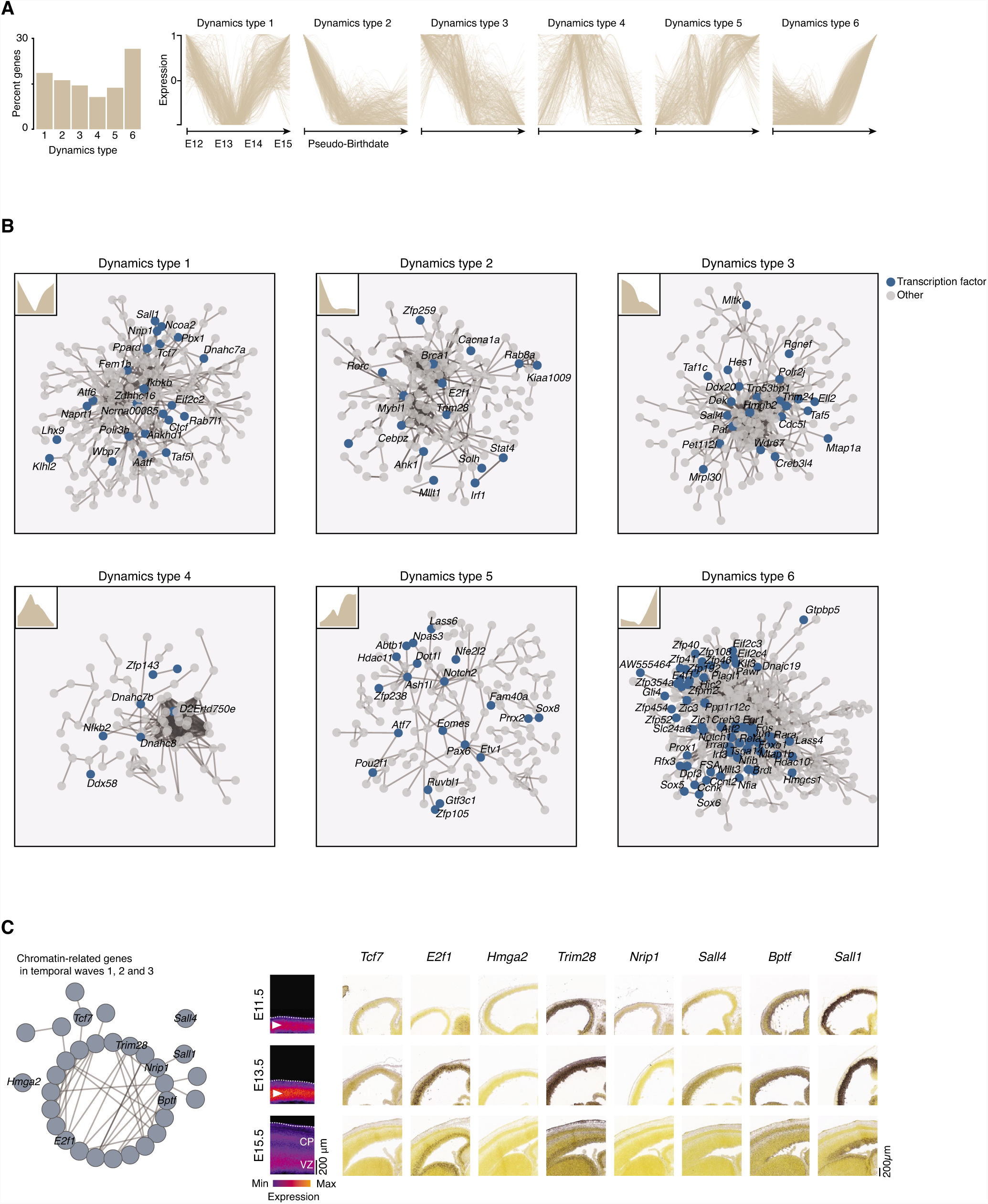
Sequential AP transcriptional states across corticogenesis. (A) Genes distribution within the six sequential AP transcriptional states. (B) Global protein-protein interactome for each AP state, from https://string-db.org/. Unassigned genes are not displayed. (C) Interactome of chromatin-related genes in AP early states. (D) In situ hybridization of chromatin-related genes at early AP states. Source of ISH: Allen Developing Mouse Brain Atlas.

**Fig. S6.**
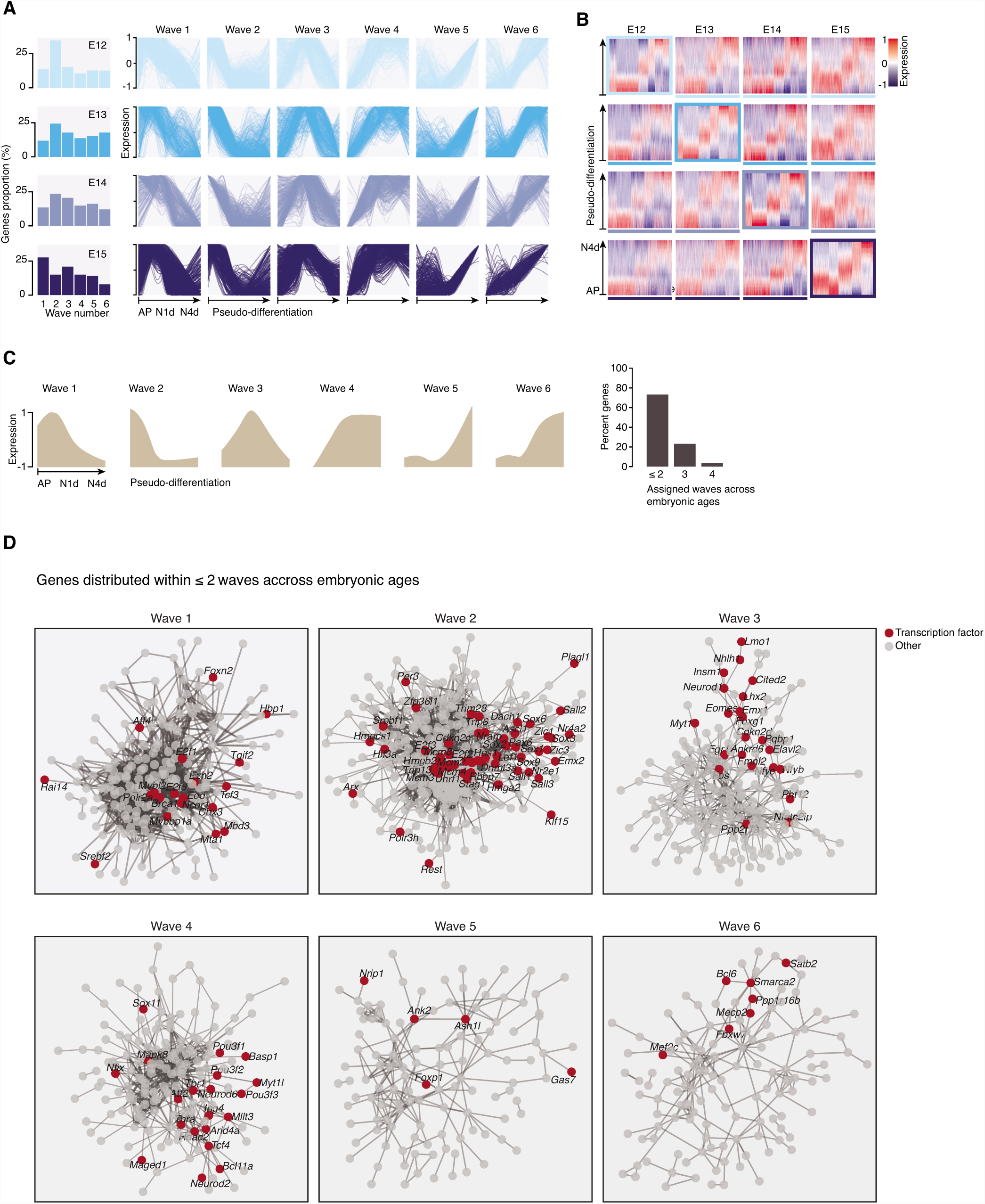
Transcriptional differentiation waves at each embryonic age. (A) Gene distribution within the six differentiation waves at each embryonic age. (B) Gene waves at each embryonic age organized depending on a reference sequence. (C) Gene distribution within waves across embryonic ages. (D) Protein-protein interactome (from https://string-db.org/) of the most stable gene within each wave. Unassigned genes are not displayed.

**Fig. S7.**
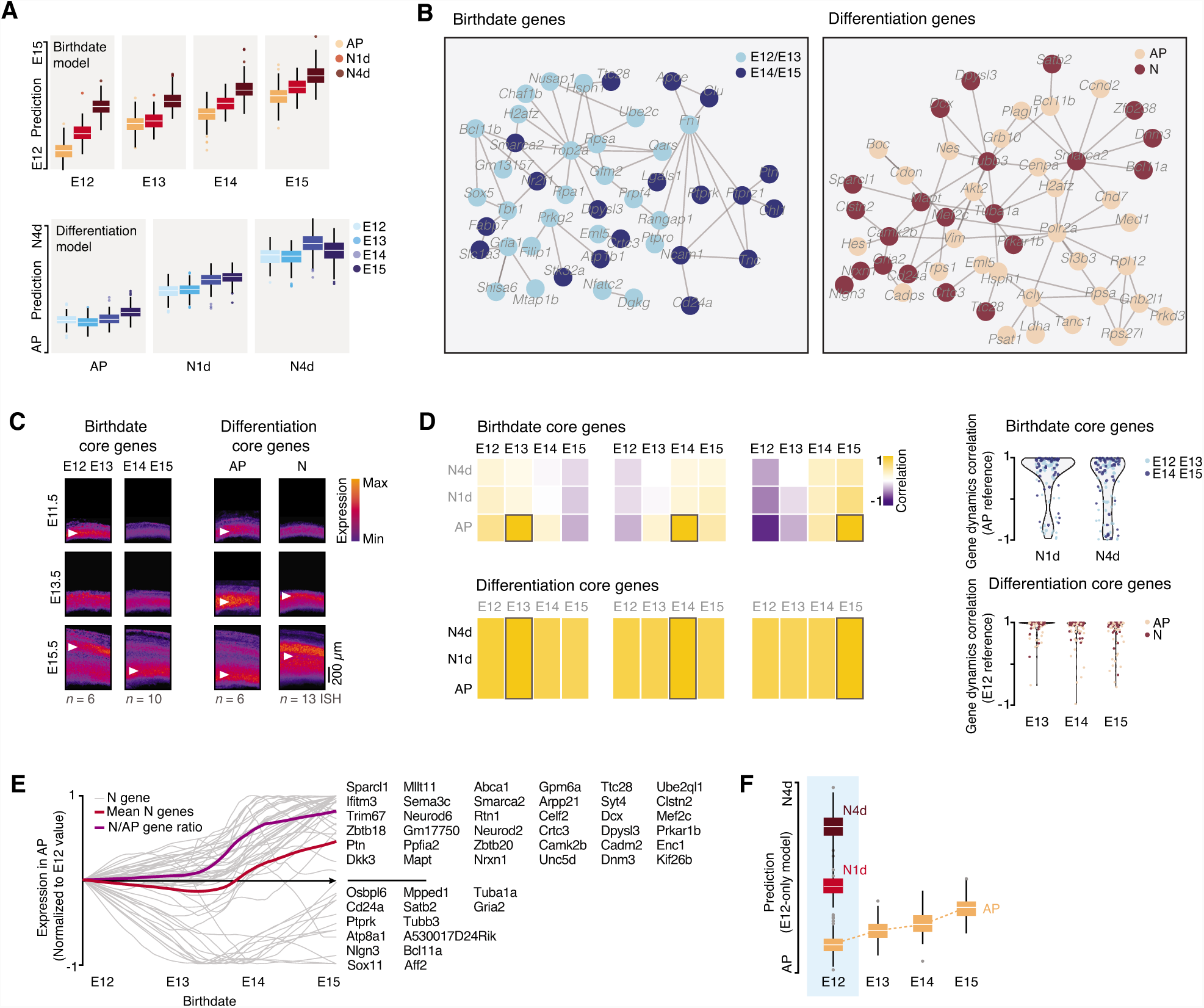
2D modelization of corticogenesis. (A) Birthdate and differentiation scores obtained from the two models for each condition. (B) Analysis of protein-protein interactions using the STRING database (http://string-db.org) suggests that gene products interact based on their temporal dynamics (left) or cellular specificity (right). Unassigned genes are not displayed. (C) Overlay of ISH from the Allen Developing Mouse Brain Atlas (www.brain-map.org) confirming the proper spatio-temporal dynamics of select core genes. Early genes: *Hes1, Hmga2, Tbr1, Fn1, Nfatc2, Sox5.* Late genes: *Nrxn1, Cttnbp2, Clu, Nr2f1, Lgals1, Bcan, Tnc, Unc5d, Slc1a3, Mfge8*. AP genes: *Cdon, Hes1, Plagl1, Nes, Hmga2, Arx*. N genes: *Trps1, Unc5d, Sox11, Nrxn, Cd24a, Mpped1, Bcl11a, Neurod6, Satb2, Dcx, Mapt, Gria2, Tubb3*. (D) Top: Birthdate-associated core genes are temporally dynamic and daughter cells acquire embryonic stage-specific transcriptional birthmarks. Bottom: In contrast, differentiation status-associated core genes are conserved across corticogenesis. Boxed area represents value of reference for correlation. Right: Correlations in gene expression dynamics stratified for early (E12, E13) and late (E14, E15) embryonic ages. (E) Expression of the core neuronal genes (n = 50) within APs increases with embryonic age. (F) E12-15 APs progressively become “neuralized”. Differentiation model build exclusively with E12 data as a training dataset; E13-E15 APs are classified as progressively more neuron-like using this model.

**Fig. S8.**
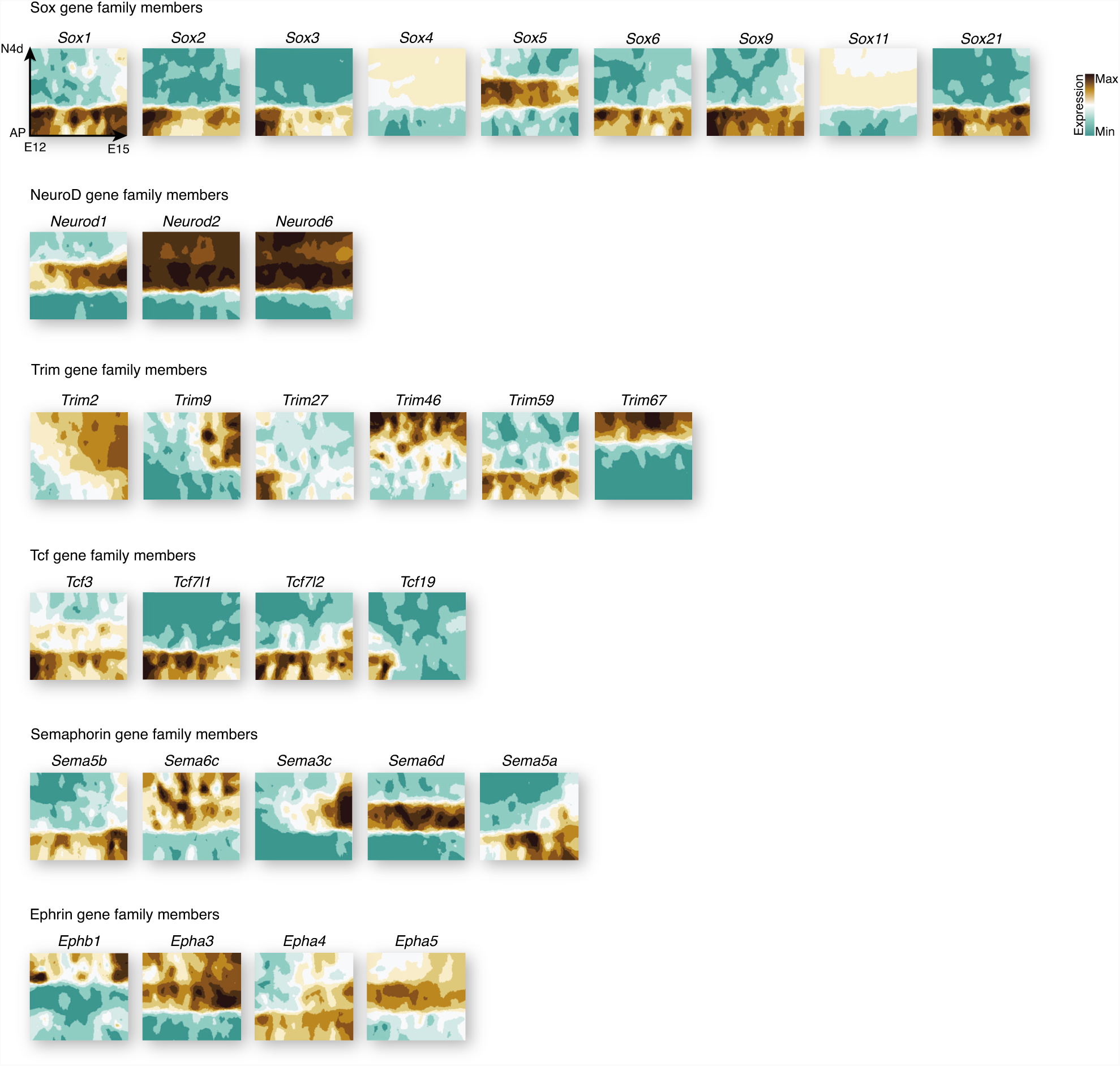
Examples of members of gene families and their associated transcriptional maps. Only the genes with the most sharply delineated expression patterns are shown for Semaphorins and Ephrins.

**Fig. S9.**
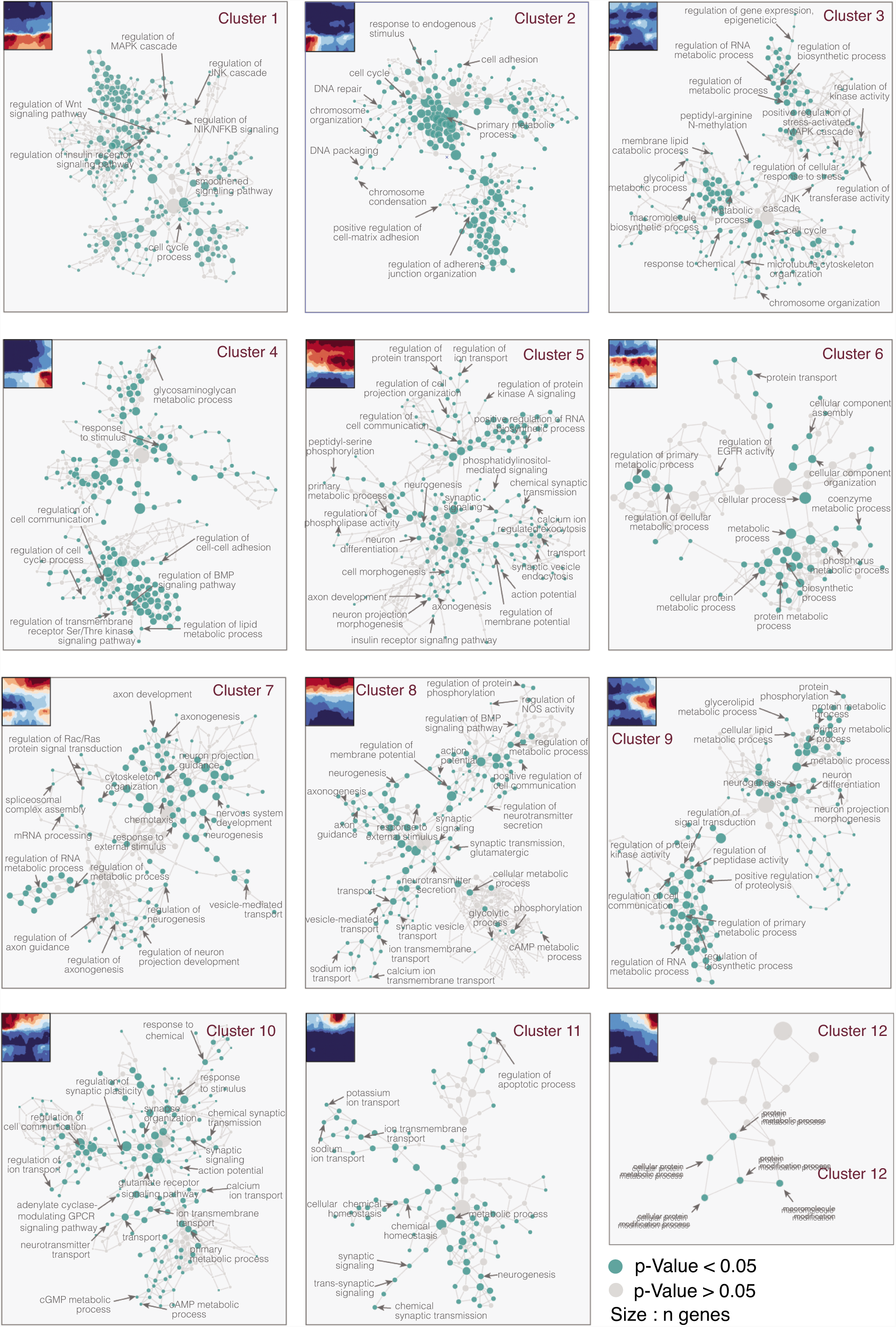
Cluster-based gene ontology networks. Display of ontological hierarchies for individual clusters highlights cluster-specific biological processes.

